# Clinically-relevant cell type cross-talk identified from a human lung tumor microenvironment interactome

**DOI:** 10.1101/637306

**Authors:** Andrew J Gentles, Angela Bik-Yu Hui, Weiguo Feng, Armon Azizi, Ramesh V. Nair, David A. Knowles, Alice Yu, Youngtae Jeong, Alborz Bejnood, Erna Forgó, Sushama Varma, Yue Xu, Amanda Kuong, Viswam S. Nair, Rob West, Matt van de Rijn, Chuong D. Hoang, Maximilian Diehn, Sylvia K. Plevritis

## Abstract

Tumors comprise a complex microenvironment of interacting malignant and stromal cell types. Much of our understanding of the tumor microenvironment comes from *in vitro* studies isolating the interactions between malignant cells and a single stromal cell type, often along a single pathway. To develop a deeper understanding of the interactions between cells within human lung tumors we performed RNA-seq profiling of flow-sorted malignant cells, endothelial cells, immune cells, fibroblasts, and bulk cells from freshly resected human primary non-small-cell lung tumors. We mapped the cell-specific differential expression of prognostically-associated secreted factors and cell surface genes, and computationally reconstructed cross-talk between these cell types to generate a novel resource we call the Lung Tumor Microenvironment Interactome (LTMI). Using this resource, we identified and validated a prognostically unfavorable influence of Gremlin-1 production by fibroblasts on proliferation of malignant lung adenocarcinoma cells. We also found a prognostically favorable association between infiltration of mast cells and less aggressive tumor cell behavior. These results illustrate the utility of the LTMI as a resource for generating hypotheses concerning tumor-microenvironment interactions that may have prognostic and therapeutic relevance.

**Summary:** RNA-seq profiling of sorted populations from primary lung cancer samples identifies prognostically relevant cross-talk between cell types in the tumor microenvironment.

## INTRODUCTION

Non-small cell lung carcinoma (NSCLC) accounts for ~80% of all lung tumors and is comprised of two major histologic subtypes: adenocarcinoma (~60%) and squamous cell carcinoma (~40%). Despite significant therapeutic efforts, overall 5-year survival for NSCLC remains a dismal 18%(*1*). While therapies that target malignant cells, such as cisplatinum-based chemotherapy and EGFR inhibitors, have led to improvements in outcomes, new therapeutic strategies are urgently needed in order to significantly improve survival of most lung cancer patients. Recent advances in tumor immunotherapy highlight the importance of targeting interactions between malignant and immune cells, with much effort focusing on the T-cell suppressive PD1/CTLA4 axes (*2*). Significant evidence points to additional complex interactions between malignant cells and other cell types comprising the tumor microenvironment. Experimental and clinical studies suggest that immune cells as well as endothelial cells and tumor-infiltrating fibroblasts play significant roles in lung cancer development and progression (*3–7*). The molecular mechanisms underlying these observations are only beginning to be understood. Rapid progress has been hindered by the reality that these cell subtypes interact in a complex network (i.e. interactome) consisting of intra- and intercellular communication via juxtacrine, autocrine and paracrine signaling. Elucidating the nature of interactions between lung cancer cells and cells comprising the tumor microenvironment could guide the development of novel therapeutic interventions.

Examples of important stromal players in NSCLC include tumor-associated macrophages (TAMs) which are a major component of the immune cell infiltrate seen in solid tumors (*8*). Macrophage-tumor cell interactions lead to release of macrophage-derived cytokines, chemokines, and growth/motility factors which in turn recruit additional inflammatory cells to the microenvironment (*9, 10*). Other immune cells commonly infiltrating lung tumors that play important roles in tumor biology include T-, B-, and NK-cells (*11–13*). Cancer associated fibroblasts (CAFs) represent another class of stromal cells that interact with the malignant cell compartment in lung cancers (*14–17*). Although several studies have found functionally important interactions between CAFs and lung cancer cells, a comprehensive understanding of their precise role in lung tumorigenesis remains lacking. These specialized fibroblasts can enhance tumor progression via multiple pathways, including synthesis of support matrices, production of promalignancy growth factors, promotion of angiogenesis, secretion of ECM proteases and pro-invasion factors such as hepatocyte growth factor, and production of immune suppressive cytokines (*15–17*).

The most common approach to studying tumor microenvironment gene expression has been to profile bulk tumors and look for cell-type-specific gene expression “clusters” in the resulting data. Interpretation becomes difficult when genes are expressed in multiple cell types. Co-expression of genes in multiple cell types within tumors occurs frequently. For example, subsets of malignant cells have been found to express genes such as vimentin and fibronectin-1 that are also expressed by fibroblasts (*18*). More recently it has become possible to perform single-cell RNAseq (scRNAseq) for hundreds to thousands of cells from a tumour sample (*19*). However the cost is still prohibitive for large cohorts, and transcriptome coverage is not complete.

Here, we dissociated primary human lung tumor samples directly after surgery, sorted individual cell subtypes based on the expression of surface markers, and performed RNA-seq analysis. We computationally identified cross-talk between different cell types in the lung tumor microenvironment, with a specific focus on prognostically-relevant associations. Through the combination of cell purification from primary tumors and gene expression profiling we have constructed a novel resource for identifying functional interactions between human lung cancer cells and their stroma: The Lung Tumor Microenvironment Interactome (LTMI; https://lungtmi.stanford.edu).

## RESULTS

Forty human primary NSCLC tumors were acquired directly from the operating room, dissociated and sorted based on CD45^+^EPCAM^-^ (pan-immune), CD31^+^CD45^-^EPCAM^-^ (endothelial cells), EPCAM^+^CD45^-^CD31^-^ (malignant cells), and CD10^+^EPCAM^-^CD45^-^CD31^-^ (fibroblasts), via our previously published flow cytometry strategy (**Figure 1a**) (*20*). We performed RNA-seq profiling on 185 samples from 36 tumors for which good quality RNA could be obtained (**Supplementary Table 1**), including unsorted bulk RNA from the majority of cases, along with six reference samples distributed between experimental batches (Stratagene Universal Reference Human RNA). This yielded an average of 48.6×10^6^ fragments per sample (range 3.8-85.4×10^6^), with mean effective mapping rate of 87.2% (range 26.6-97.1%). Out of a total of 28034 expressed protein-coding genes across all cell types, 5790 (21%) were expressed in all four and 9918 (35%) were expressed in only one (**Supplementary Figure 1**), indicating that a large fraction of genes are expressed in specific TMI subpopulations. All patients were treatment-naïve, and clinical characteristics of the cohort are shown in **Supplementary Table 2**. Sample SNP profiles were compared to verify identities (**Supplementary Figure 2**) (*21*). After data normalization and summarization of expression at the gene level, we performed batch correction to remove differences between flow cells and observed that this generally improved concordance between transcriptomes from replicates (n=11; **Supplementary Figure 3**). Multidimensional scaling (MDS) analysis of the 1000 most variable genes across sorted populations showed separation of the malignant, fibroblast, immune, and endothelial cells (**Figure 1b**). There was no separation between individual populations isolated from adenocarcinoma versus squamous cell carcinoma by MDS. We next performed unsupervised hierarchical clustering analysis on the same 1000 genes and again observed clear separation of profiles from the different sorted cell types (**Figure 1c**). With one exception (T29 CD31^+^), replicates were immediately adjacent to each other in the sample-wise dendrogram.

**Figure 1.**
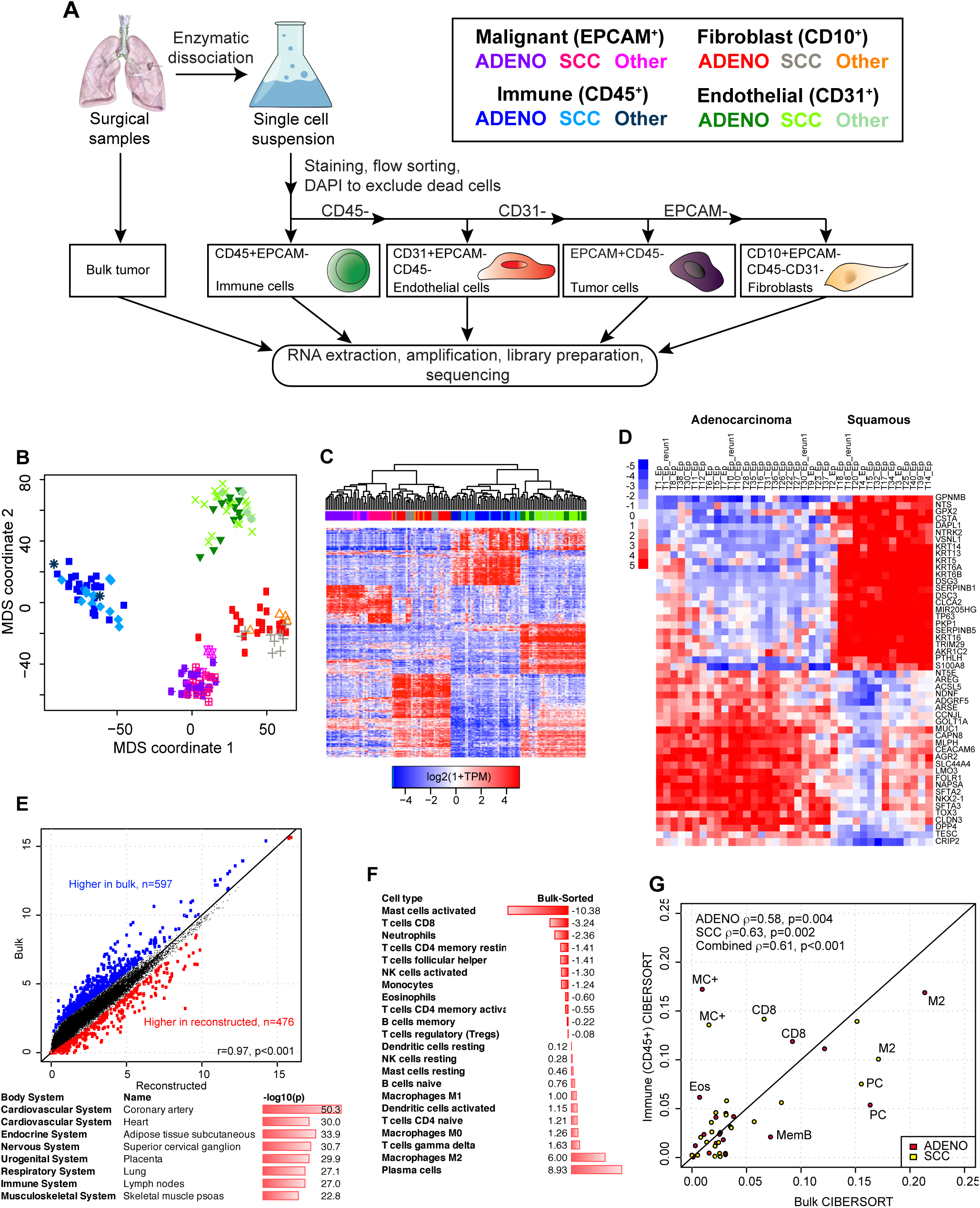
**(a)** Schema for dissociation, flow-sorting, and RNA-seq profiling. **(b)** Multidimensional scaling analysis of transcriptomes of cell types sorted from surgically resected primary human NSCLC tumors. Axis units are arbitrary. Cell types are depicted by colors as in 1a. **(c)** Unbiased hierarchical clustering of sorted samples. **(d)** Top 25 most differentially expressed genes between malignant cells from adenocarcinoma and SCC. **(e)** Comparison of bulk vs reconstituted transcriptomic profiles. Shown are average values across all samples for each gene measured by RNA-seq. Panel below shows functional enrichment of genes higher in bulk for tissues that were not sorted for profiling. **(f)** Average percentage difference in immune cell types deconvolved in bulk vs sorted CD45+ populations showing enrichment of activated mast cell profiles by sorting, and conversely loss of plasma cells. **(g)** CIBERSORT deconvolution of immune populations in adenocarcinoma (pink) and SCC (light blue) identifies similarities and differences immune cell types that are relatively depleted (below diagonal) or enriched (above diagonal) by dissociation and sorting. MC+ = activated mast cells; PC = plasma cells; M2 = M2-polarized macrophages; MemB = memory B-cells; CD8 = CD8 T-cells; Eos = Eosinophils.

Within the malignant population, adenocarcinomas clustered apart from SCC as expected. The one tumor in our cohort that was called NSCLC NOS (Not Otherwise Specified) based on histopathology clustered with adenocarcinoma, whereas three other tumors T9 (adenosquamous), T23 (fetal), and T37 (invasive mucinous adenocarcinoma) clustered with SCC. In supervised analysis, 1168 genes were differentially expression between malignant cells from adenocarcinoma versus SCC tumors (local FDR <1%) with 931 being more highly expressed in adenocarcinoma, and 237 being lower (**Supplementary Table 3**). Distinguishing genes included classic basal keratins (KRT5, KRT6A, KRT6B, KRT13, KRT14) that along with TP63 were more highly expressed in SCC (**Figure 1d**). Conversely, NKX2-1, and mucins (MUC1, MUC5B) were more highly expressed in adenocarcinoma, as were ROS1 and CLDN3.

One potential limitation of experimental strategies that involve dissociation and sorting of cells is that these procedures could distort their transcriptomes prior to RNA-seq profiling. To permit analysis of this phenomenon, we also performed RNA-seq on bulk tissues that were frozen immediately after surgical dissection (**Supplementary Table 1**). We then compared the “ground truth” bulk transcriptional profile of each sample to the computationally reconstructed one defined by combining the profiles of individual populations weighted according to their relative abundance in the tumor. The latter was defined by deconvolving the bulk transcriptomes using CIBERSORT (*22*), with the sorted sample transcriptomes used to construct a signature matrix (**Supplementary Figure 4 and Supplementary Tables 4 and 5**). Reconstructed profiles largely recapitulated bulk profiles (R=0.97, **Figure 1e**, and **Supplementary Table 6**). We further explored these differences by ranking all genes by their difference between bulk and reconstructed profiles, and compared these to an atlas of body tissues using the Nextbio Correlation Engine(*23*). Off-diagonal genes higher in bulk were ones archetypally expressed on cell populations that were not isolated by sorting, including muscle- and nerve-related genes (**Figure 1e**, **Supplementary Table 6**). To further isolate the effects of dissociation and sorting, we applied CIBERSORT to the bulk tumors and the sorted immune samples using our previously validated signature matrix of 22 immune cell types (LM22; (*22*)) and compared deconvolution results. Relative proportions of infiltrating leukocytes were similar across adenocarcinoma and SCC in both our RNA-seq data and previously published microarray studies (**Supplementary Figure 5**). Direct comparison of deconvolution results based on bulk vs sorted immune cells showed that some immune subtypes had higher or lower inferred proportions in sorted immune cells, suggesting that they were more sensitive to dissociation and/or sorting (**Figure 1f,g)**. These included lower than expected levels of plasma cells in immune sorted populations; and higher levels of activated mast cells and eosinophils. Plasma cells are systematically lost during flow sorting, whereas activation/degranulation of mast cells might be triggered by sorting. Taken together, these results suggest that our experimental strategy left the transcriptomes of most populations largely intact, but identify specific populations that are sensitive to flow sorting. These findings are likely also relevant for scRNA-seq analyses.

### The LTMI reveals a complex transcriptional landscape of secreted ligands and their receptors across NSCLC tumor sub-populations

To identify avenues for cross-talk between cell types in adenocarcinoma and SCC, we integrated the LTMI data with the FANTOM5 resource of ligand-receptor interactions, and the PRECOG resource of prognostic associations between bulk gene expression and overall survival (**Figure 2a**) (*24, 25*). We examined the potential complexity of cell-cell interactions by comparing the number of populations in which a ligand was expressed with the number of populations in which its cognate receptor was expressed, using TPM>10 as a threshold, to be consistent with the criteria used by FANTOM5 (**Figure 2b and Supplementary Table 7**). In both NSCLC histologies, the most frequent pattern was many-to-one, where all four sorted populations expressed a ligand, but only one population expressed its receptor; however, this pattern was not statistically significantly more prevalent than others (p=0.11 and p=0.18 respectively in adenocarcinoma and SCC by chi-squared test). In general, both ligands and receptors could be seen to be uniquely or ubiquitously expressed, suggesting that transcriptional regulation of cellular crosstalk is occurring at the level of both ligand and receptor activity.

**Figure 2.**
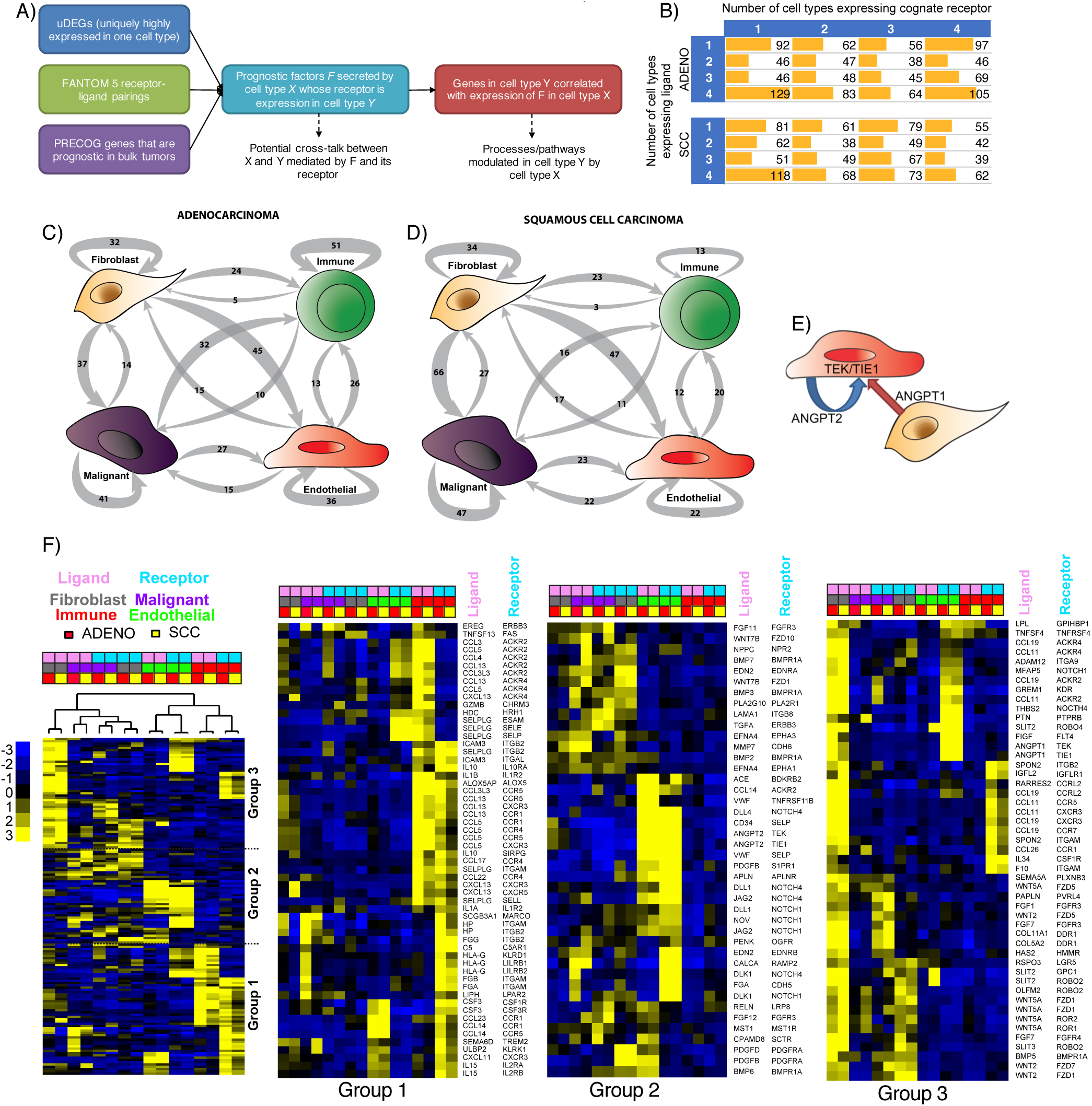
**(a)** The Lung Tumor Microenvironment Interactome (LTMI) integrates data generated in this study, the FANTOM5 resource of ligand-receptor pairs, and PRECOG for prognostic associations of genes in bulk tumor samples. **(b)** Potential complexity of inter-cell-type signaling via secreted factors. Ligands or receptors were defined as significantly expressed in a cell type if they had TPM>10, as in the FANTOM5 study. **(c, d)** Potential cross-talk between cell types in adenocarcinoma (c) and squamous cell carcinoma (d). Shown are the number of ligand-receptor pairs where each is a uniquely differentially expressed gene (uDEG) in the indicated cell-type. Arrows X->Y indicate that the ligand is a uDEG in cell type X, while the corresponding receptor is a uDEG in cell type Y. **(e)** Expression patterns of ligands and receptors pair that are highly expressed (TPM>10) in single cell types (corresponding to the 1-1 entries for adenocarcinoma and SCC in panel (b). Pink indicates ligand whereas blue indicates receptor. **(f)** Expanded view of the three groups shown in panel (e). **(g)** ANGPT1 and ANGPT2 compete antagonistically for receptor binding and have opposite prognostic associations in NSCLC. They are expressed on fibroblasts and endothelial cells respectively, with expression of their known receptors being predominantly in endothelial cells.

We identified genes that were significantly differentially expressed in specific cell types relative to others, separately in adenocarcinoma and SCC, focusing on those that were over-expressed in a single cell type relative to all others i.e. uniquely differentially expressed genes (uDEGs), at FDR <1% with a minimum 2-fold difference in expression (Methods). We intersected these with genes coding for putative secreted factors and cell surface proteins. Expression levels of these genes are frequently associated with survival outcomes in lung cancer, however the cell type producing these factors is often unknown (**Supplementary Table 8**). In both adenocarcinoma and SCC, the most prolific expressors of uDEG ligands were fibroblasts (**Figure 2c,d and Supplementary Table 9**). Their corresponding uDEG receptors were most commonly expressed by endothelial and malignant cells (**Figure 2c,d**). Malignant cells also highly expressed many ligands, but their receptors were most frequently also expressed in malignant cells, suggesting autocrine signaling. In adenocarcinoma, receptor uDEGs for ligands differentially expressed in immune cells were most often also on immune cells; with a similar pattern seen in endothelial cells. In contrast, in SCC we did not observe a bias towards expression of both ligand and receptor within immune and endothelial populations.

We noted that the vascular growth signaling angiopoietin genes ANGPT1 and ANGPT2 were expressed by fibroblasts and endothelial cells respectively whereas the cognate receptors encoded by TEK and TIE1 were only expressed on endothelial cells. In bulk tumors, high ANGPT1 expression is associated with good overall survival while high ANGPT2 is associated with poor survival. Their products function as a rheostat competing for receptor binding, and the LTMI suggests that this occurs in an intra-cell type fashion (**Figure 2e)**. We further examined the pattern of expression of ligand-receptor pairs where each was highly expressed in a single cell type (**Figure 2f**). One major group (Group 3) showed a pattern where fibroblasts were prolific expressors of ligands whose receptors were expressed on every possible cell type, suggesting autocrine and paracrine signaling. This included BMP (bone morphogenic protein) signaling pathways involving BMP3 and BMP2 which promote cell growth. Group 2 displayed a prominent enrichment for NOTCH-related signaling within the endothelial compartments of both adenocarcinoma and SCC. An enrichment for immune compartment expression of ligands/receptors dominated Group 1, with potential autocrine and paracrine cross-talk potential. The latter was mainly with endothelial cells (in both adenocarcinoma and SCC), or malignant cells (in adenocarcinoma only).

Overall, our results indicated potential for highly complex inter-population communication via ligand-receptor signaling, particularly initiating from fibroblasts to endothelial or malignant cells, and autocrine influences within the malignant compartment.

### Identification of clinically-relevant cell type cross-talk using the LTMI

We sought to identify and validate cross-talk between cell types in the LTMI that had potential clinical relevance (**Figure 2a**). To this end, we focused on genes that were associated with patient survival and that were expressed in a specific cell type as assessed by RNA-seq. Among the resulting potential interactions, we selected two (GREM1 and TPSAB1, encoding mast cell tryptase MCT) for experimental validation. These represented cross-talk between fibroblasts and malignant cells (GREM1); and between immune and malignant cells (TPSAB1).

### High expression of Gremlin-1 by fibroblasts stimulates proliferation of lung adenocarcinoma tumor cells

High expression of GREM1, encoding for the secreted factor Gremlin-1, is associated with poor overall survival in lung adenocarcinoma but not squamous cell carcinoma (PRECOG meta-Z: +4.11 in adenocarcinoma vs −0.75 in SCC). Our data showed it to be expressed strongly in fibroblasts from both histologies, but not in other cell-types (**Figure 3a**). GREM1 inhibits bone morphogenetic protein (BMP) signaling by binding BMP ligands and preventing their interaction with their receptors(*26*). Additionally, GREM1 has been shown to bind and activate the vascular endothelial growth factor (VEGF) receptor Kinase Insert Domain Receptor (KDR, also known as VEGFR2, one of two receptors for VEGF), which is expressed in endothelial cells of both adenocarcinoma and SCC. Interestingly, KDR is also expressed in the malignant compartment in adenocarcinoma at 3-fold higher levels than in SCC (p=1×10^−5^, t-test; see also Figure 2f).

**Figure 3.**
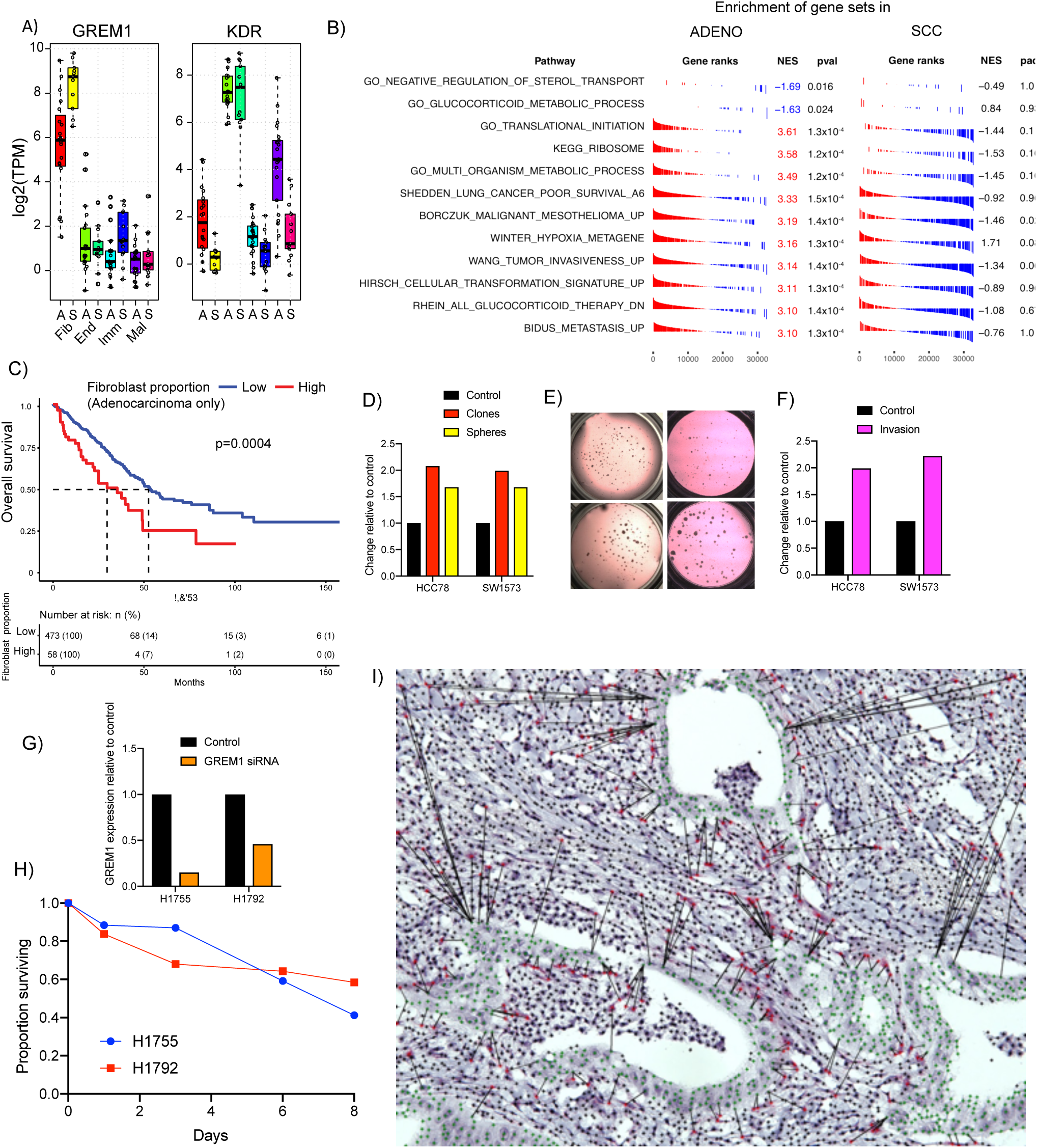
**(a)** GREM1 (encoding the secreted factor Gremlin-1) is highly expressed on fibroblasts in adenocarcinoma and SCC. Its receptor KDR is highly expressed in endothelial cells of both adenocarcinoma and SCC, and also in malignant cells from adenocarcinoma but not SCC. **(b)** Expression of GREM1 in fibroblasts is positively correlated with expression of proliferation and invasiveness related genes in malignant cells in adenocarcinoma (all adjusted p-values <0.05), but not in SCC. **(c)** High levels of fibroblasts inferred in adenocarcinoma from TCGA are associated with less favorable overall survival. **(d,e,f)** Treatment of low GREM1-expressing adenocarcinoma cell lines HCC78 and SW1573 with recombinant Gremlin-1 protein resulted in increased number of clones (red), sphere formation in 3-D culture (yellow), and invasion as evaluated by *in vitro* trans-well migration assays (magenta). **(g)** si-RNA knockdown resulted in decreased GREM1 expression in both H1755 and H1792 adenocarcinoma cell lines, which normally express it highly. **(h)** Knockdown of GREM1 expression reduced survival in both cell lines that highly express it. **(i)** Representative stain for GREM1 RNA (ref) shows expression confined to fibroblasts, that spatially colocate preferentially with leading edge of malignant cell nests. Malignant cells are highlighted in green. Black bars show closest malignant cell to each GREM1+ fibroblast.

We next sought evidence for a role for GREM1 in cross-talk between fibroblasts and malignant cells by using the LTMI to correlate gene expression levels in malignant cells from adenocarcinoma with the level of GREM1 in fibroblasts from the same tumors. Expression levels of genes involved in translation initiation, ribosomal biogenesis and invasiveness in malignant cells were positively correlated with GREM1 expression in fibroblasts from the same patient in adenocarcinoma but not in SCC (**Figure 3b; see also Supplementary Table 10**). Genes related to cellular transformation and hypoxia were also higher when GREM1 was higher in adenocarcinoma, but not SCC. Additionally, higher adenocarcinoma fibroblast GREM1 correlated with lower malignant cell glucocorticoid metabolism gene expression. Together, these observations suggested that GREM1 production by fibroblasts might induce a more aggressive malignant cell behavior in adenocarcinoma but not squamous cell carcinoma. To further test this, we evaluated the relationship between fibroblast content and overall survival in TCGA adenocarcinoma and SCC tumors with CIBERSORT using the signature matrix defined by our purified cell populations (**Supplementary Table 5**). Patients with a higher inferred proportion of fibroblasts had worse overall survival in adenocarcinoma (p=0.01 as a continuous variable, likelihood ratio test) but not in SCC (p=0.83, not shown). An optimal dichotomization of adenocarcinoma into patients with fibroblast proportion higher or lower than 17% robustly separated survival curves (p=0.0004, log-rank test; **Figure 3c**).

Based on the results from the LTMI, we sought to functionally test if GREM1 can alter behavior of lung cancer cells. Lung cancer cell lines express GREM1 at varying levels, with ~5500-fold range across SCC lines and nearly 13,000-fold across adenocarcinomas as measured in the Cancer Cell Line Encyclopedia (**Supplementary Table 11**) (*27*). To test a positive causal association of GREM1 with malignant cell behavior we treated adenocarcinoma cell lines with low intrinsic expression of the gene (HCC78 and SW1573) with recombinant GREM1. Treatment with GREM1 increased both 2D colony and 3D tumorsphere formation by approximately 2 fold (**Figure 3d,e**). Additionally, GREM1 treatment resulted in significantly higher migratory potential using *in vitro* trans-well migration assays (**Figure 3f)**. Thus, exogenous GREM1 increases aggressiveness of lung cancer cells *in vitro*.

As noted above, some lung cancer cell lines express high levels of GREM1, suggesting a potential tumor-promoting autocrine role in a subset of lung cancers. Consistent with this we observed a range of GREM1 expression in the malignant cells from human tumors with a small number of outliers expressing significant levels of GREM1 (**Figure 3a**). To test if GREM1 may have an autocrine function in these cells, we knocked down the transcript in high GREM1 expressing H1755 (which does not express the KDR receptor) and H1792 (which does express KDR) adenocarcinoma cells using siRNA. Knock-down reduced GREM1 transcript levels by 85% in H1755 and 54% in H1792 (**Figure 3g**), and reduced survival of both cell lines by up to 50% after 8 days (**Figure 3h)**.

### GREM1-expressing fibroblasts are preferentially spatially located adjacent to malignant cells

We further verified that GREM1 expression was confined to fibroblasts using In-Situ RNA hybridization on tumor tissues (**Figure 3i**). Interestingly, visual inspection indicated that fibroblasts expressing GREM1 clustered around nests of cancer cells, suggesting a potential juxtacrine interaction between these cell types mediated by this pathway. In order to assess the spatial distribution of GREM1 positive cells we stained and digitally imaged four tissue samples from tumors corresponding to very low, low, medium or high GREM1 expression. We developed an automated image processing pipeline (Methods) to detect nuclei and classify cells as GREM1 positive vs. negative. We used this pipeline to quantitatively evaluate our qualitative observation that GREM1+ cells tend to be physically closer to tumor cells than other stromal cells. We manually annotated tumor regions in the four images and calculated the distance to the nearest tumor cell for every stromal cell, both GREM1+ and GREM1-. We then compared the distribution of these distances for GREM1+ vs. GREM1-using a Mann-Whitney U test for difference in the mean. For all three samples with GREM1 expression the GREM1+ cells were significantly closer on average to malignant cells than GREM1-cells (p=3×10^−16^, 1×10^−7^ and 1×10^−10^ for the low, medium and high tissue samples respectively – no GREM1+ cells were detected in the very low GREM1 expression image). To further confirm this result we performed a simulation study, repeatedly resampling the stroma nuclei as being GREM1+ vs. GREM1-while maintaining the same number of GREM1+ cells. We used the median distance of the GREM1+ cells to the nearest tumor cells as a test statistic, *T*. For all three samples with GREM1 expression, out of 10^5^ simulations, *T* was never as small as for the observed configuration, implying a p-value of <1×10^−5^ in each case.

### Levels of infiltrating mast cells negatively correlate with tumor proliferation in adenocarcinoma and SCC

To further demonstrate of the utility of the LTMI, we investigated potential associations between the immune and malignant compartments. We noted that TPSAB1 (Tryptase Alpha/Beta 1) was highly expressed in sorted immune cells from both adenocarcinoma and SCC (p<2.2×10^−16^ by ANOVA; **Figure 4a**), and is favorably prognostic in both histologies across multiple datasets in PRECOG(*24*). Among a panel of 22 different immune cell types TPSAB1 expression was nearly 30-fold more highly expressed on mast cells (**Supplementary Figure 6**). We performed cross-population enrichment analysis by ranking genes in malignant cells by their correlation to TPSAB1 expression in immune cells across the cohort (**Supplementary Table 12**). In adenocarcinoma, high TPSAB1 expression in immune cells correlated with reduced malignant cell expression of proliferation and cell cycle genes as well as of genes related to metastasis (**Figure 4b**). Few gene sets positively correlated with TPSAB1 expression, but included olfactory receptor genes and genes down-regulated in gefitinib-resistant NSCLC. In SCC, we again observed negative association of immune TPSAB1 expression with metastasis and extracellular matrix genes in malignant cells, as well as VEGF and EGF signaling pathway genes (**Figure 4c**). However, interestingly proliferation genes in SCC malignant cells positively correlated with immune TPSAB1, in contrast to adenocarcinoma.

**Figure 4.**
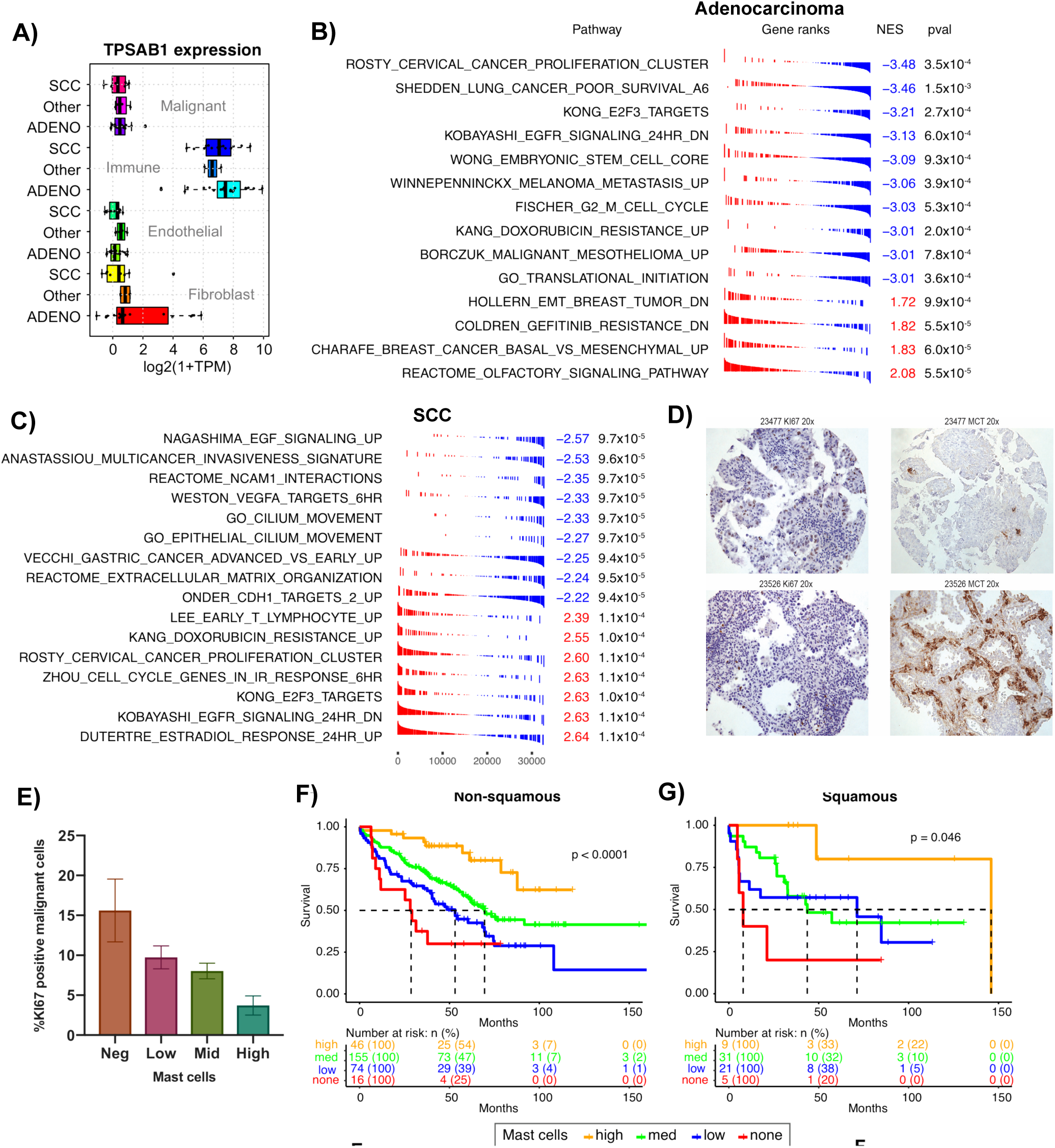
**(a)** TPSAB1 (encoding Tryptase α/β 1) is highly expressed in immune cells in both adenocarcinoma and SCC. **(b,c)** TPSAB1 expression in immune cells was negatively associated with proliferation and metastasis-related genes in adenocarcinoma (b); while in SCC there was a negative association with invasiveness and angiogenesis but a positive association with proliferation. **(d)** Representative stains for cellular proliferation marker KI67, and MCT in samples that had high (top) and low (bottom) expression of TPSAB1. Shown are 20X magnification image; see Supplementary Figures 8 and 9 for 40X and 60X. **(e)** Primary adenocarcinomas with higher numbers of infiltrating mast cells had a lower proportion of KI67-positive (proliferating) malignant cells (*p*=0.003; F-test). **(f, g)** High numbers of mast cells in both primary adenocarcinomas (f) and SCC (g), assessed by tissue microarray staining for mast cell tryptase (MCT), were associated with better overall survival. Mast cell counts were assigned to pre-defined “none”, “low”, “medium”, and “high” categories by pathologist.

We validated the prognostic relevance of mast cells in NSCLC by immunohistochemical (IHC) staining of a lung tumor tissue microarray (TMA) for MCT (mast cell tryptase, encoded by TPSAB1). The lung TMA (n=389 samples) was stained for MCT, and each core was scored for the number of mast cells by a pathologist. Mast cell infiltration was similar across NSCLC histologies, but higher in adenocarcinoma in situ relative to other types (**Supplementary Figure 7**). Within adenocarcinoma, mast cell counts were significantly higher in Stage 1 vs Stage 3 (p=0.006 by t-test) but not in Stage 1 vs Stage 2 or Stage 2 vs Stage 3 (**Supplementary Figure 7**). There was no difference in mast cell levels across stages of SCC, though the modest sample size (n=66) limited the statistical power of this analysis.

Mast cell counts were converted to levels of “High”, “Intermediate”, “Low”, and “Negative” (Methods). In order to validate the relationship of mast cell levels to tumor proliferation the same TMA was stained for the proliferation marker KI67 (**Figure 4d**, see also **Supplementary Figures 8** and **9**). The proportion of KI67-positive malignant adenocarcinoma cells was lower in tumors with high vs low/intermediate numbers of mast cells (**Figure 4e**; p=0.003, ANOVA F-test), consistent with the gene set based analysis of our sorted RNA-seq data. Negative-, low-, and intermediate-levels of mast cells all conferred worse overall survival than high mast-cell levels whether considered across only non-squamous NSCLC (n=214, **Figure 4f**) or SCC (n=66, **Figure 4g**). Multivariable analysis indicated that mast cell levels carried prognostic information independent of stage (**Supplementary Table 13**), and Kaplan-Meier analysis within stages I, II and III separately confirmed that the level of mast cell infiltration was prognostic across NSCLC and within adenocarcinoma (**Supplementary Figure 10**).

### The LTMI: a resource for exploring lung tumor microenvironment interactions

To facilitate investigation of relationships between transcriptional profiles within the lung TMI we developed an online resource, the Lung Tumor Microenvironment Interactome (https://lungtmi.stanford.edu). Users can select from sets of genes that are prognostic, expressed in a specific population, and/or encode for secreted or surface factors. Alternatively, a list of genes of specific interest can be entered manually. Given this set of genes, the LTMI interface can display differential expression between different sub-populations in adenocarcinoma and SCC. Correlations can be extracted between expression levels of these genes in a cell type of interest compared to other cell types. Gene set enrichment analysis, as performed in this study, can be applied to the resulting correlative output. A tutorial in the resource is available to recapitulate the results described here relating GREM1 in fibroblasts to malignant cell transcriptional programs.

## DISCUSSION

As with other malignancies, most research efforts on lung cancer have focused on the transformed cells themselves. This has led to the identification of important pathways and individual genes involved in oncogenesis such as EGFR, KRAS, and ALK (*28–31*). Significantly less attention has been directed at investigating possible contributions of the tumor microenvironment to cancer formation, progression and treatment response, although this is a burgeoning area of interest. Here we developed a unique resource by profiling human primary lung tumors that were dissociated and sorted directly after surgical resection.

Prior applications of computationally-derived regulatory networks have used whole tumor high-throughput data to gain insight into mechanisms underlying hematological cancers (*32–36*). More recently, such computational approaches have been extended to solid tumors (*37–39*). Previous work on profiling the tumor microenvironment has often been accomplished through the use of laser capture microdissection (LCM) in a variety of tumors, including those of the breast and lung (*40, 41*). However, it is difficult to separate endothelial cells, fibroblasts, and infiltrating immune cells using LCM and these are therefore usually lumped as one stromal sample in such studies. Our approach for gene expression profiling malignant and stromal cells within primary tumors involves dissociating the tumor tissues and then purifying individual cell subtypes based on the expression of surface markers.

By RNA-seq profiling of cell types from lung tumors we found that GREM1, high levels of which are associated with worse patient outcomes, is specifically expressed on fibroblasts in the adenocarcinoma microenvironment. The LTMI identified a positive association between fibroblast GREM1 expression and malignant cell proliferation genes. Gremlin-1 has been shown previously to induce proliferation of normal lung cells, and to be over-expressed in adenocarcinoma, but not SCC, compared to normal lung. However to the best of our knowledge neither its fibroblast origin, nor a specific role in stimulating proliferation of lung cancer lines has been noted. Cancer associated fibroblasts (CAFs) represent a major class of stromal cells that interact with the malignant cell compartment in lung cancers (*14*). CAFs appear biologically distinct from fibroblasts present in benign microenvironments (*42*). Although several studies have found functionally important interactions between CAFs and lung cancer cells (*43–45*), the role of Gremlin-1 identified using the LTMI appears to be novel. Adenocarcinoma cell lines express GREM1 variably. Si-RNA knockdown in high-expressing cell lines resulted in reduced proliferation independent of KDR receptor expression. However, our data suggest that in primary tumors, receptor expression is required. Interfering with this TME interaction may therefore represent a novel therapeutic opportunity.

Interactions between malignant cells and infiltrating immune cells are another major class of microenvironmental interactions within lung tumors. There have been conflicting reports concerning a role for mast cells in cancer, and specifically in lung tumors, with some finding them to be a favorable prognostic factor, and others an adverse factor [refs]. Using the LTMI we identified and validated a favorable prognostic association of mast cell infiltration in lung tumors. This finding was consistent with a novel inverse correlation between mast cell infiltration and numbers of KI67 positive malignant cells. The mechanistic influence of mast cells on NSCLC malignant cells requires further investigation, however they have been proposed to have cytolytic activity in breast cancer(*46*)

Limitations of our study include the focus on four pre-defined sub-populations, the potential impact of cell dissociation and sorting on transcriptional profiles, and the restriction to expression data. Nonetheless, we were able to identify and validate associations between malignant and stromal cell types. In the future, single-cell RNA-seq will increasingly be used to dissect the tumor microenvironment and will allow further resolution of transcriptional properties of malignant and stromal sub-populations within the TMI. However it is not yet practical for large cohorts, and still has technical limitations that preclude full coverage of the transcriptome.

In conclusion, we have developed a publicly available resource called the Lung Tumor Microenvironment Interactome that allows interrogation of potential interactions between cell subpopulations within human lung tumors. We anticipate that this resource will complement scRNA-seq analyses, and facilitate future studies of lung cancer biology that will allow identification of novel drug targets for improving treatment outcomes for this devastating disease.

## MATERIALS AND METHODS

All patient samples in this study were collected with informed consent for research use, and approved by Stanford Institutional Review Board in accordance with the Declaration of Helsinki. Freshly resected surgical tumor samples from patients with NSCLC were dissociated and sorted as described (*20*) using A700 anti-human CD45 clone HI30 (pan-leukocyte cell marker), PE anti-human CD31 clone XWM59 (endothelial cell marker), APC anti-human EpCAM clone X9C4 (epithelial cell marker), and PE-Cy7 anti-human CD10 clone XHI10a (fibroblast marker). All antibodies were obtained from BioLegend (San Diego, CA). Library preparation and sequencing were performed as described previously.

### Cell lines and reagents

The human lung cancer cell lines were obtained from American Type Culture Collection. All cell lines were cultured in RPMI-1640, supplemented with 10% FBS and 100 mg/L penicillin/streptomycin, and maintained at 37C with 5% CO^2^.

### Effect of gremlin-1 in lung adenocarcinoma cells

Recombinant Grem-1 protein (500ng/ml) was added to lung adenocarcinoma cell lines (HCC78, SW1573) with low intrinsic GREM1 expression in 2D or 3D culture. The clones were stained with crystal violet and enumerated at 10-14 days after seeding the cells. Effect of Grem-1 on lung adenocarcinoma cell migration and invasion were evaluated using *in vitro* trans-well migration assays. Recombinant Grem-1 protein (500ng/ml) was added to lung HCC78 and SW1573 cell lines in 3D culture. Numbers of spheres were counted 8-10 days after seeding the cells on matrigel.

### si-RNA knockdown of GREM1, viability and clonogenic assays

siRNA (30M) targeted against Grem-1 (siGrem-1) was used to decrease GREM1 mRNA expression in lung adenocarcinoma cell lines with high GREM1 expression (H1755 and H1792). Viability of si-Grem-1 transfected cells was examined using the CellTiter 96 Non-Radioactive Cell Proliferation Assay (MTS), according to the manufacturer’s protocol (Promega BioSciences). For clonogenic assays (2D), identical number of cells with or without treatment were reseeded at low density in 6-well plates in triplicate and incubated at 37C under 5%CO_2_. After 10 to 12 days, plates were washed, fixed in 50% methanol, and stained with 0.1% crystal violet and then the number of colonies was counted. Evaluation of colony formation was also conducted in 3D cell culture using matrigel (Corning) and cell culture inserts for 24-well plates (Corning). After 10 to 12 days, the number of spheres were enumerated under a light microscope.

### In vitro migration and invasion assays

Effect of Grem-1 in the invasion of cells were assayed using the BD BioCoat Matrigel Invasion Chambers (BD Bioscience). Each well of a 24-well plate contained an insert with an 8-mm pore size PET (polyethylene terephthalate) membrane. Inserts coated with a thin layer of Matrigel basement membrane matrix were used to measure the ability of the cells to invade through the reconstituted basement membrane. 1X10^5^ cells were seeded inside the insert with medium containing 1% serum. High serum (10%) medium was then added to the bottom chamber of 24-well plates to serve as a chemoattractant. After 24 hours, the membranes were washed, stained, then separated with a sterile scalpel and mounted on a glass slide. The number of migrating cells were counted under a light microscope.

### Western blot analysis

Total protein extracts were harvested from cell lines and prepared for immunoblotting. Membranes were probed with rabbit monoclonal antibodies (Cell Signaling Technology) including anti-Phospho-Smad1(Ser463/465)/Smad5 (Ser Ser463/465)/Smad9 (Ser465/467) (D5B10), anti-c-Myc (D84C12) and anti–β-actin (D6A8), followed by secondary antibodies conjugated to horseradish peroxidase. β-actin protein levels were used as loading controls. Western blots were quantified with the Adobe Photoshop Pixel Quantification Plug-In (Richard Rosenman Advertising & Design).

### Quantitative PCR

qRT-PCR analysis was utilized to analyze expression changes of gremlin-1, c-myc, p21 and GAPDH. Total RNA was isolated from cells using the Paris Kit (Ambion). One microgram of total RNA was reverse transcribed The High Capacity cDNA Reverse Transcription Kit (Applied Biosystems) as specified by the manufacturer. qRT-PCR was done using SYBRGreen PCR Master Mix (Applied Biosystems) and an ABI PRISM 7900 Sequence Detection System (Applied Bio-systems). Primers for PCR amplifications (Supplementary Table S1) were designed using Primer 3 Input (version 0.4.0). Relative mRNA levels were calculated using the 2 ΔΔ C_t_ method.

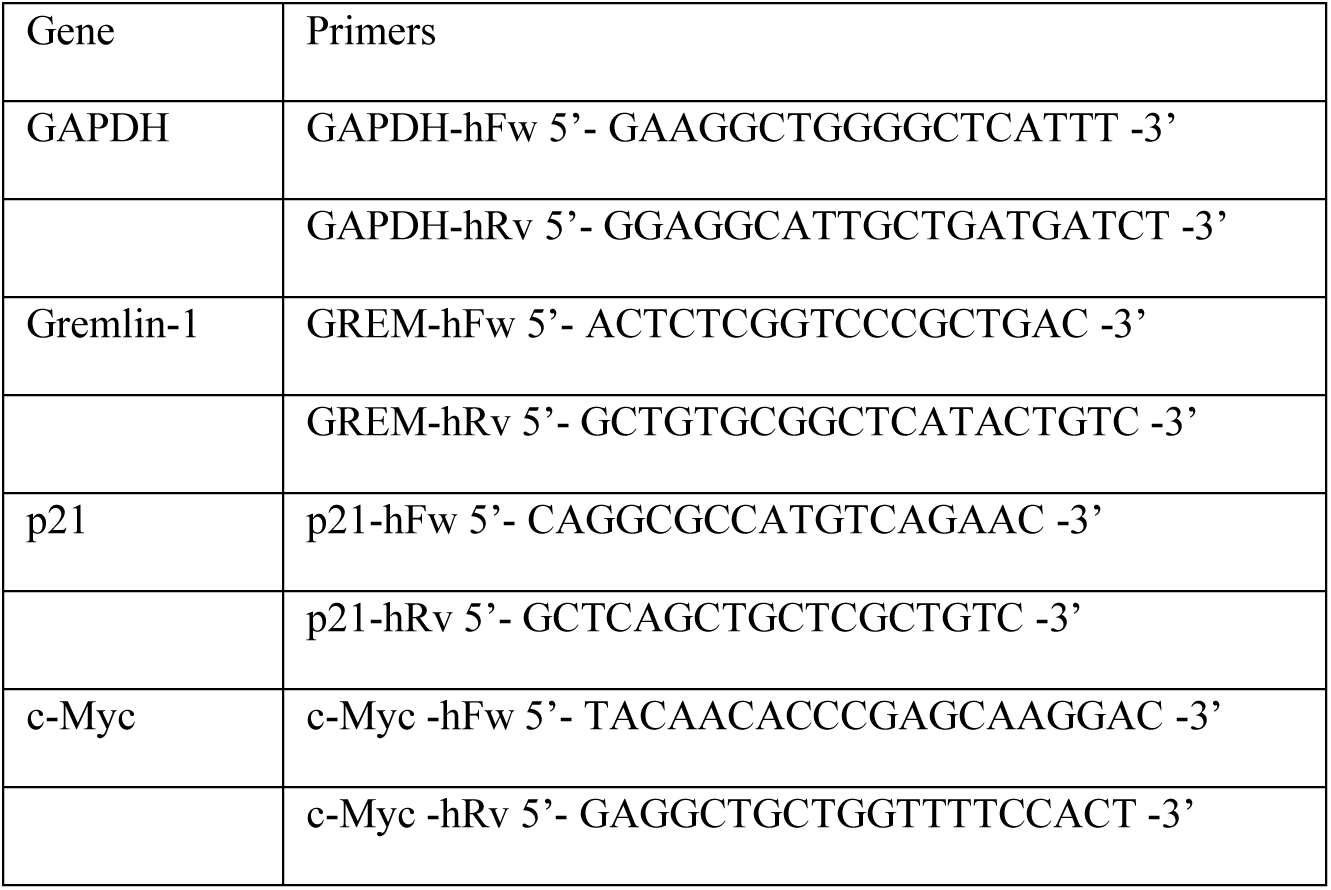

### Tissue Microarray staining and analysis

The tissue microarray (TMA) was cut into 4µm thick sections, deparaffinized and hydrated. For the Ki-67 staining, the TMA slide was subjected to Epitope Retrieval Solution 2 (ER2, Leica) antigen retrieval and stained with a prediluted anti-Ki-67 antibody (Mouse, Clone MIB-1, Dako #M7240) using an automated immunostainer. For the MCT (mast cell tryptase; corresponding to TPSAB1 gene) staining, the TMA slide was subjected to Cell Conditioning 1 (CC1, Ventana) antigen retrieval and stained with a prediluted anti-MCT antibody (Mouse, Clone G3, Millipore #MAB1222) using an automated immunostainer. Mast cell counts were assigned to pre-defined categories by pathologists as follows: “none” when 0 mast cells, “low” when 1-9 mast cells, “medium” when 10-30 mast cells, and “high” when greater than 31 mast cells were present in each entire 0.6 mm core.

### Computational Analysis

Briefly, paired-end reads were aligned to the human genome (GRCh38) using STAR version 2.5.0 (*47*), with Gencode v23 transcriptome annotation (*48*) using a 2-pass approach. Alignment files were de-duplicated, and further processed using the Genome Analysis Toolkit (GATK). Expression levels were also quantified using Salmon v0.4.2 (*49*). Individual Salmon runs were integrated into a single expression matrix using the *tximport* package in R, using TPM (transcripts per million) to summarize to the gene level (*50*). Despite the use of the Nugen kit for eliminating ribosomal rRNA, in common with previous studies (*51*), we found that a large and variable proportion of reads derived from mitochondrial rRNA, specifically MT-RNR1 (Mitochondrially Encoded 12S RNA) and MT-RNR2 (Mitochondrially Encoded 16S RNA). Accordingly, we renormalized the data matrix by removing these transcripts and rescaling each sample to have TPM sum to 10^6^. This preserved the relative ranking within each sample but rescales between samples, eliminating the distorting effect of the two mitochondrial rRNAs without changing the relative ranking on genes in TPM space. For subsequent analysis, we eliminated genes that had mean TPM<1 in all sample subtypes (Bulk, fibroblast, endothelial, immune, malignant) in all histologies (adenocarcinoma, SCC, or “other”). For clustering, visualization and subsequent analyses we used the moderated log of TPM i.e. log_2_(1+TPM).

We observed that there were significant batch effects between sequencing lanes based on Salmon quantification of RNA levels, with the majority of transcripts being significantly associated with sequencing lane. We applied batch correction at the level of flowcell identity using ComBat (*52*), with histology/sub-population as a model matrix, to avoid eradicating biologically meaningful signals. After this step, the lane-level batch effect was largely eliminated. RNA-seq data are available in the Gene Expression Omnibus under accession number GSE111907. The lung-TMI website interface (http://lungtmi.stanford.edu) was built using R/Shiny. In order to assess the spatial distribution of GREM1 positive cells we immuno-stained and digitally imaged four tissue samples corresponding to very low, low, medium or high GREM1 expression. We developed an automated image processing pipeline using the Pillow 2.7.0 fork of the Python Imaging Library to detect nuclei and classify GREM1 positive vs. negative. This pipeline involved:

1. Non-negative Matrix Factorization (using the NMF function from scikit-learn) to separate the GREM1 and hematoxylin channels.
2. Applying a Laplacian filter with radius 6 to the hematoxylin channel followed by non-maximum suppression to detect nuclei centers.
3. Applying a Gaussian filter with radius 6 to the GREM1 channel and evaluating at the nuclei centers, followed by thresholding at 0.1 to detect positive vs. negative GREM1 expression.

### Verification of sample identities

In order to verify that there were no sample swaps during preparation or sequencing, we performed pairwise comparison of all BAM files using bam-matcher (*21*). This tool uses a set of known common SNPs and computes the overlap between genotypes of samples. We used the Freebayes genotyping option, a depth threshold of 10 for considering a position, and the largest available set of common SNPs (n=7550).

### Secreted and surface factors

We compiled lists of genes encoding potential secreted and surface factors from several sources. For secreted factors we included: known chemokines and cytokines obtained by searching Entrez gene; genes whose SwissPROT function or localization included “secreted” as a keyword; and an additional list of WNT- and Sonic Hedgehog genes that we noted were not included among the previous groups. For surface factor genes, we took the computationally inferred list previously described as the “Surfaceome” (*53*). We also incorporated information on ligand-receptor pairs defined by the FANTOM5 consortium.

## ACKNOWLEDGEMENTS

This work was supported by grants from the National Institutes of Health/National Cancer Institute (NIH/NCI) U01 CA154969 (SKP, MD). The Genome Sequencing Service Center provided by Stanford Center for Genomics and Personalized Medicine Sequencing Center is supported by NIH grant S10OD02014. We thank Alex Dobin for advice and input regarding filtering and selecting splice junctions in two-pass alignments with STAR (*47*). The Redcap clinical databased used in this project is supported by grant UL1 TR001085 from NIH/NCRR.

## SUPPLEMENTARY TABLE LEGENDS AND SUPPLEMENTARY FIGURES

**Supplementary Table 1**

Histology of patient tumor samples, and populations processed for RNA-seq. Crosses indicate “not done”, usually due to low cell numbers, or poor RNA quality. Samples with double ticks were sequenced twice.

**Supplementary Table 2**

Clinical characteristics of the cohorts used in this study (gene expression and Tissue Microarray)

**Supplementary Table 3**

Differentially expressed genes between sorted malignant populations from adenocarcinoma and squamous cell carcinoma

**Supplementary Table 4**

CIBERSORT signature matrix derived from RNA-seq of sorted populations from adenocarcinoma and SCC. Separate signatures were used for malignant cells, pan-NSCLC for fibroblasts, endothelial cells, and immune cells.

**Supplementary Table 5**

CIBERSORT deconvolution outputs for U01 signature matrix versus sorted and bulk populations. P-values are for the overall deconvolution; see Newman et al for full discussion.

**Supplementary Table 6**

Differences between average gene expression across transcriptome in bulk vs. reconstructed profiles. Relevant to Figure 1e.

**Supplementary Table 7**

Number of populations that highly express ligands and their receptors in adenocarcinoma and SCC. A cut-off of TPM>10 was used for comparison with the original FANTOM5 study.

**Supplementary Table 8**

PRECOG prognostic z-scores for genes that encode secreted or surface proteins in adenocarcinoma and SCC.

**Supplementary Table 9**

Number of uDEG ligands and receptors (Data underlying Figure 2c and 2d). Shown are the number of cases where a ligand (rows) is a uDEG in the indicated population and its cognate receptor is a uDEG in a population (column). Top panel: adenocarcinoma; bottom panel: SCC.

**Supplementary Table 10**

Gene Set Enrichment analysis applied to ranked gene lists representing correlation between expression levels in malignant sorted populations vs GREM1 expression in fibroblasts. The c2 (curated gene sets) and c5 (Gene Ontology) gene sets were used from the Molecular Signatures Database. Selected enrichment results are shown in Figure 3b).

**Supplementary Table 11**

Expression of GREM1 in cell lines from the Cancer Cell Line Encyclopedia. The values shown are the log2 of Affymetrix signal intensity which has a range of 0 to 15. Two adenocarcinomas with low GREM1 (HCC78 and SW1573 SCC), and two with high GREM1 (H1755 and H1792) were selected for experimental validations based on availability.

**Supplementary Table 12**

Gene Set Enrichment analysis applied to ranked gene lists representing correlation between expression levels in malignant sorted populations vs TPSAB1 (mast cell tryptase) expression in the pan-immune population. The c2 (curated gene sets) and c5 (Gene Ontology) gene sets were used from the Molecular Signatures Database. Selected enrichment results are shown in Figures 4b,c).

**Supplementary Table 13**

Gene Set Enrichment analysis applied to ranked gene lists representing correlation between expression levels in malignant sorted populations vs GREM1 expression in fibroblasts. The c2 (curated gene sets) and c5 (Gene Ontology) gene sets were used from the Molecular Signatures Database. Selected enrichment results are shown in Figure 3b).

**Supplementary Figure 1:**
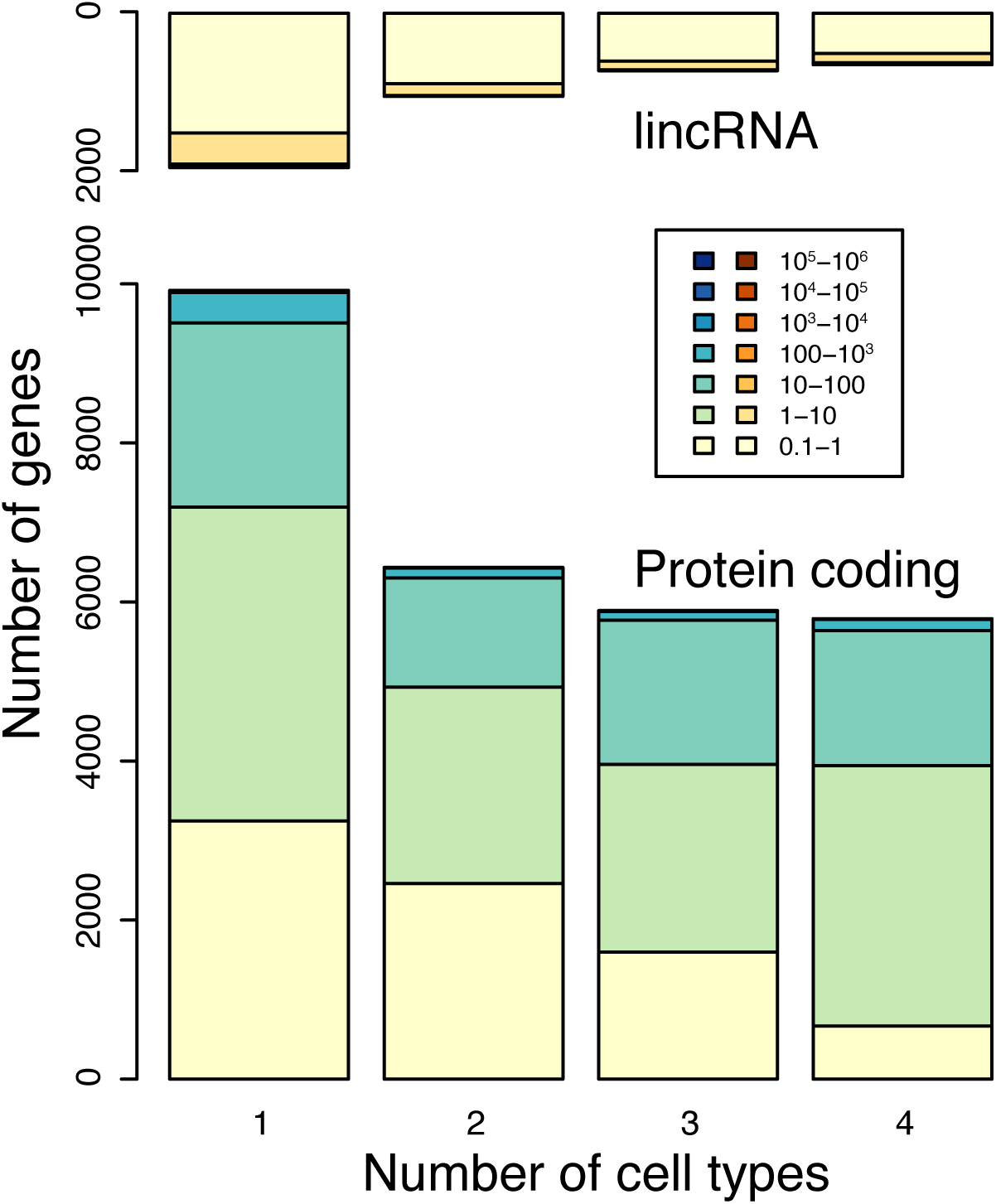
Transcriptional output across feature types and tissues. Summary of the number of protein coding genes and lincRNAs that are expressed at specific TPM thresholds in 1, 2, 3, or 4 sorted populations.

**Supplementary Figure 2.**
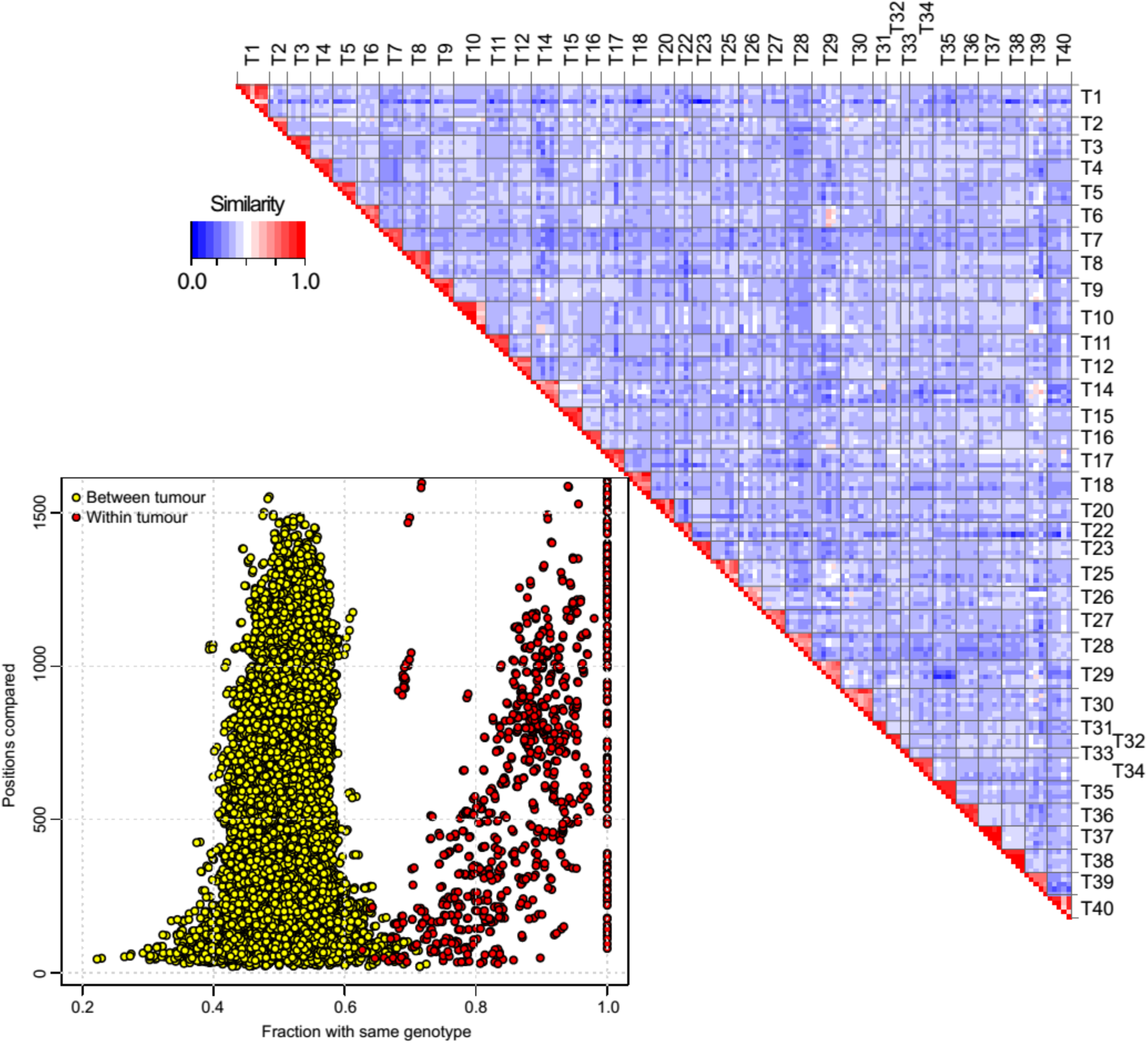
Check of sample identities by comparing SNPs. All pairwise comparisons were done using *Bammatch*. Similarity was computed as the proportion of shared SNPs (upper right of figure). Bottom left panel shows sample similarity vs number of SNPs compared. Red are samples from the same tumor; yellow from different tumors.

**Supplementary Figure 3.**
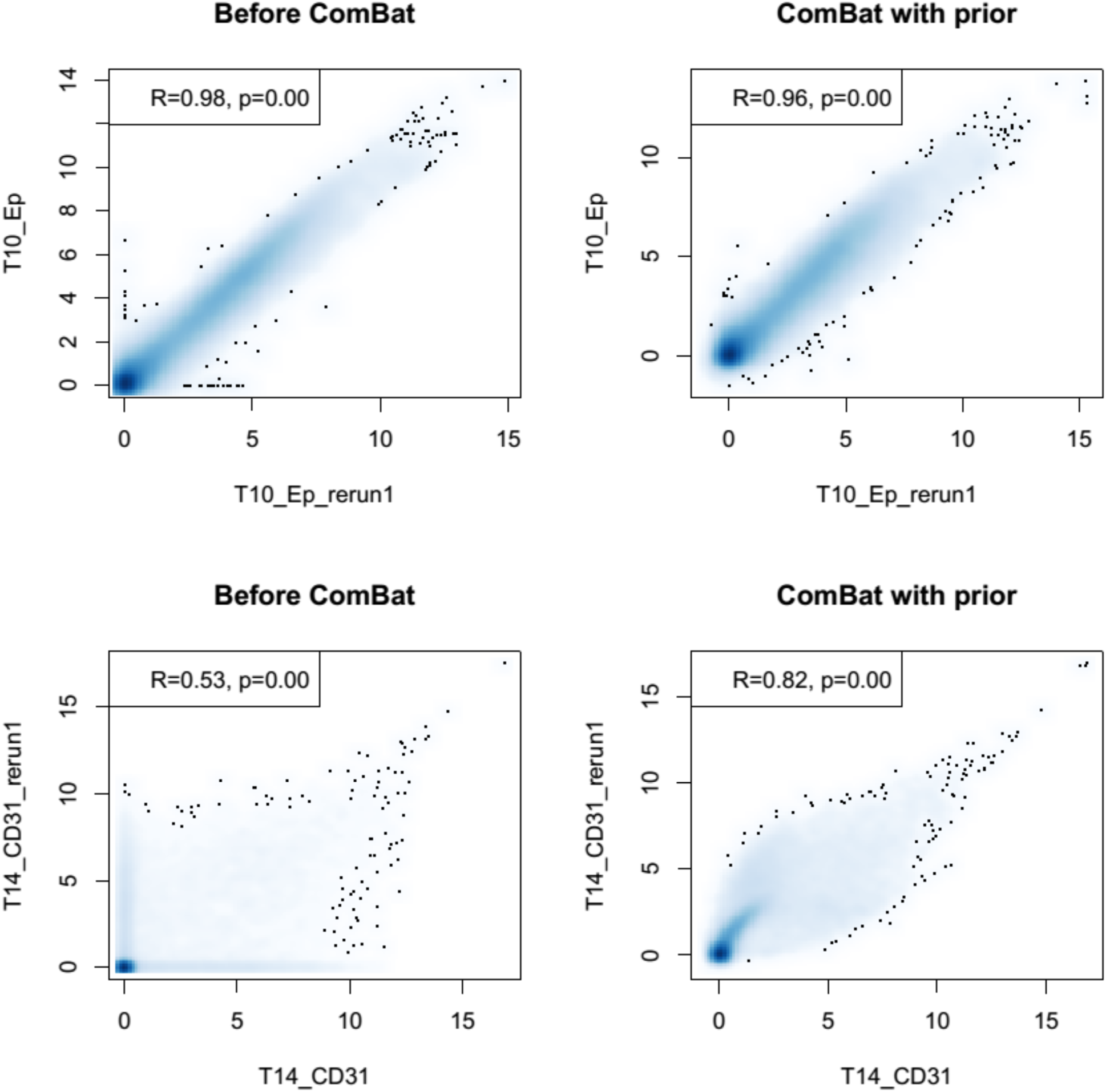
Example of effects of batch correction on two representative samples; one with good initial concordance and one with poor concordance. The blue density plot represent all genes quantified; dots are outliers.

**Supplementary Figure 4.**
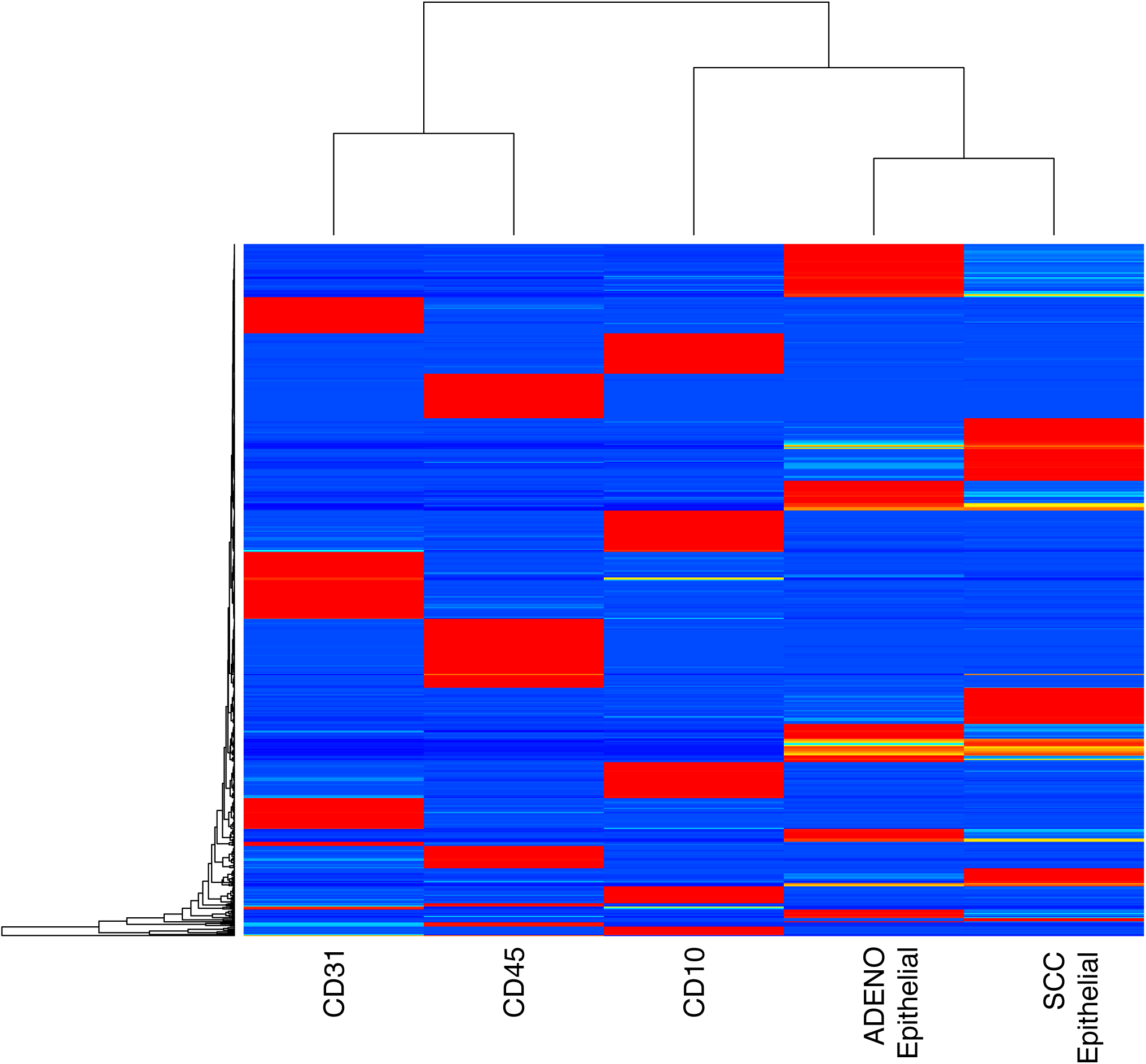
CIBERSORT signature matrix. Depiction of CIBERSORT signature matrix used for inferring proportions of cell types in bulk tumors. Rows represent genes (n=652); see Supplementary Table 4 for the actual genes and their average expression in each cell type class.

**Supplementary Figure 5.**
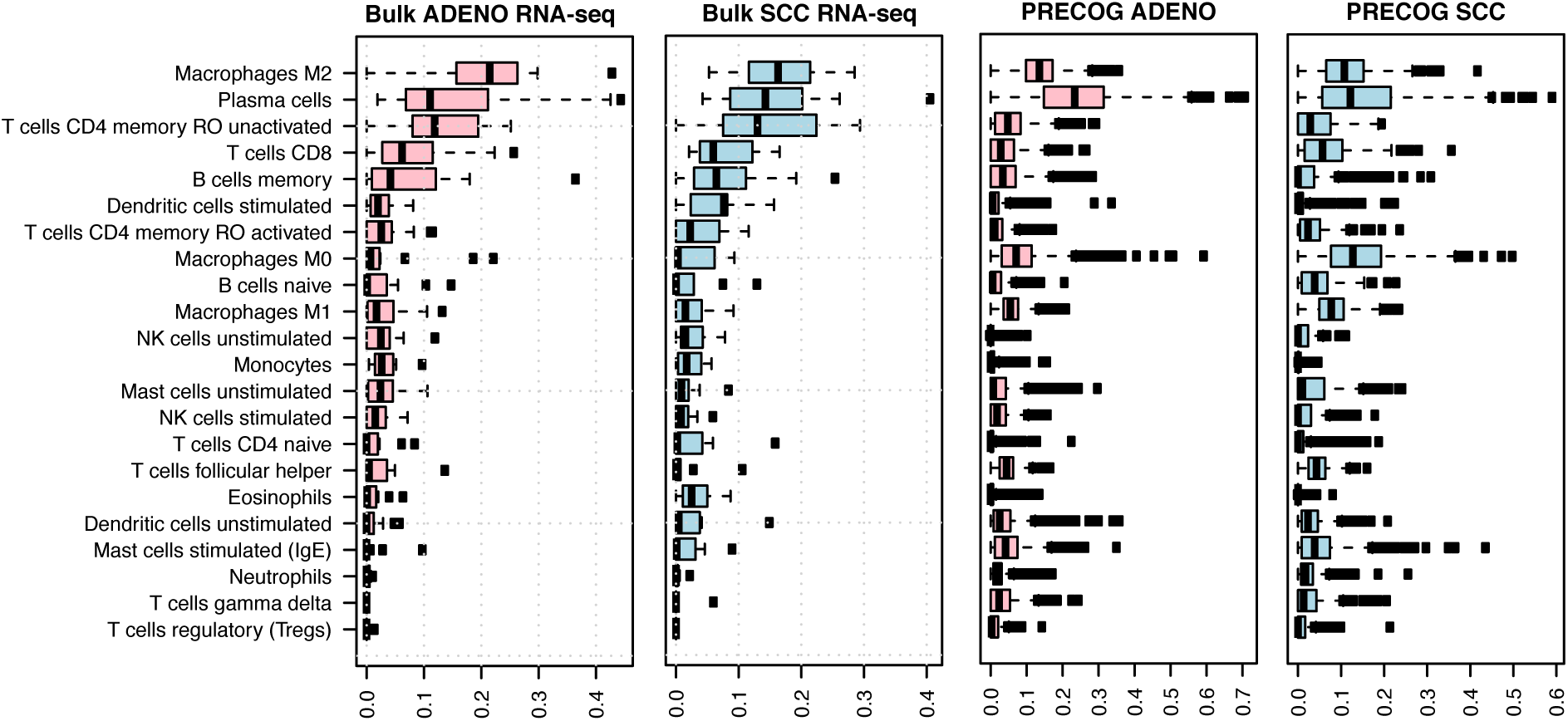
CIBERSORT deconvolution of immune populations in bulk RNA-seq and PRE-COG microarray samples. Comparison of immune proportions inferred in lung adenocarinoma and SCC by CIBERSORT from bulk RNA-seq (this study) and from microarrays (PRECOG)

**Supplementary Figure 6.**
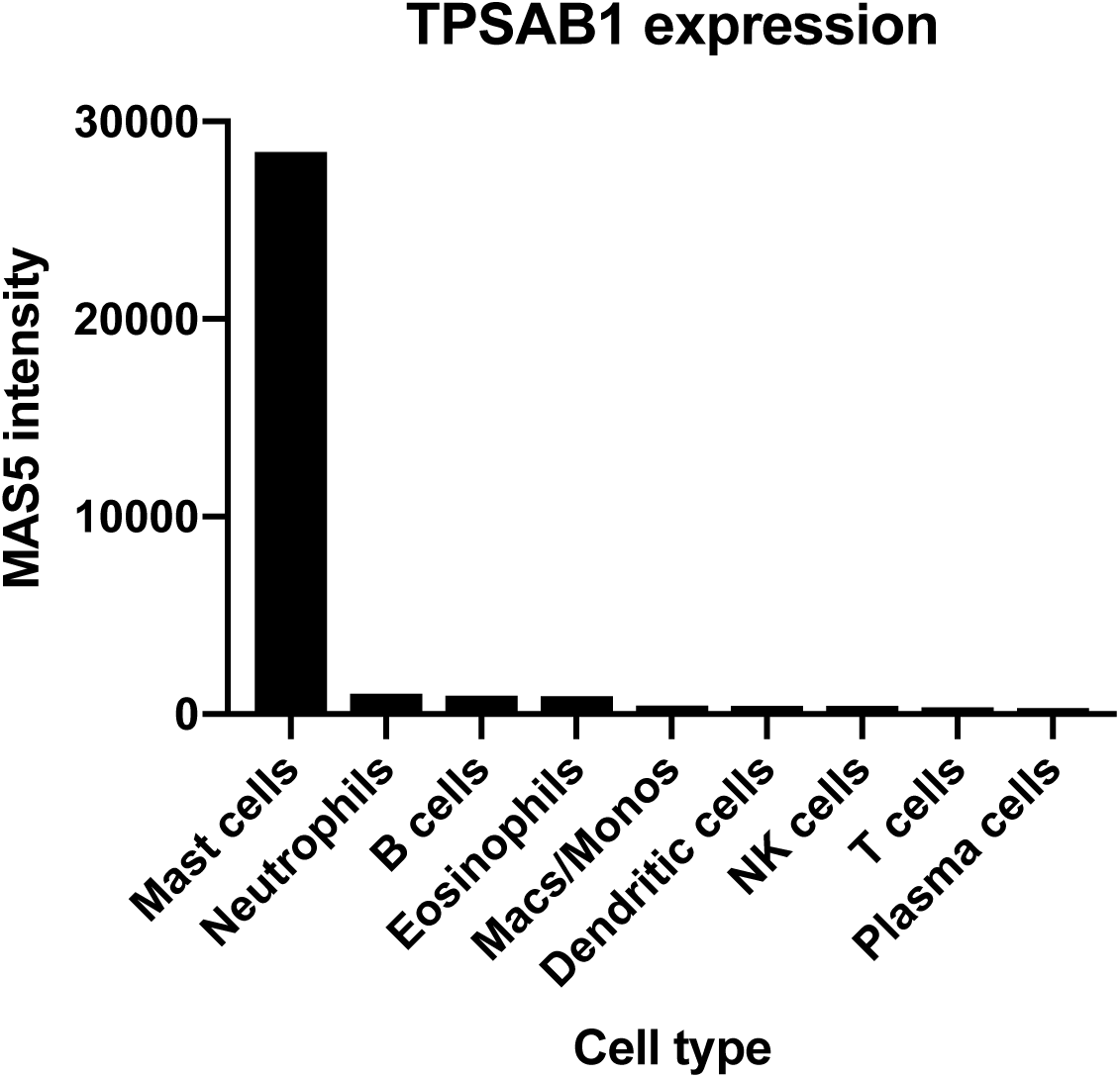
TPSAB1 expression is 30-fold higher in mast cells relative to other immune cell types in the LM22 signature matrix from CIBERSORT. Bars show the Affymetrix MAS5 intensity levels, averaged across replicates for the indicated cell types.

**Supplementary Figure 7:**
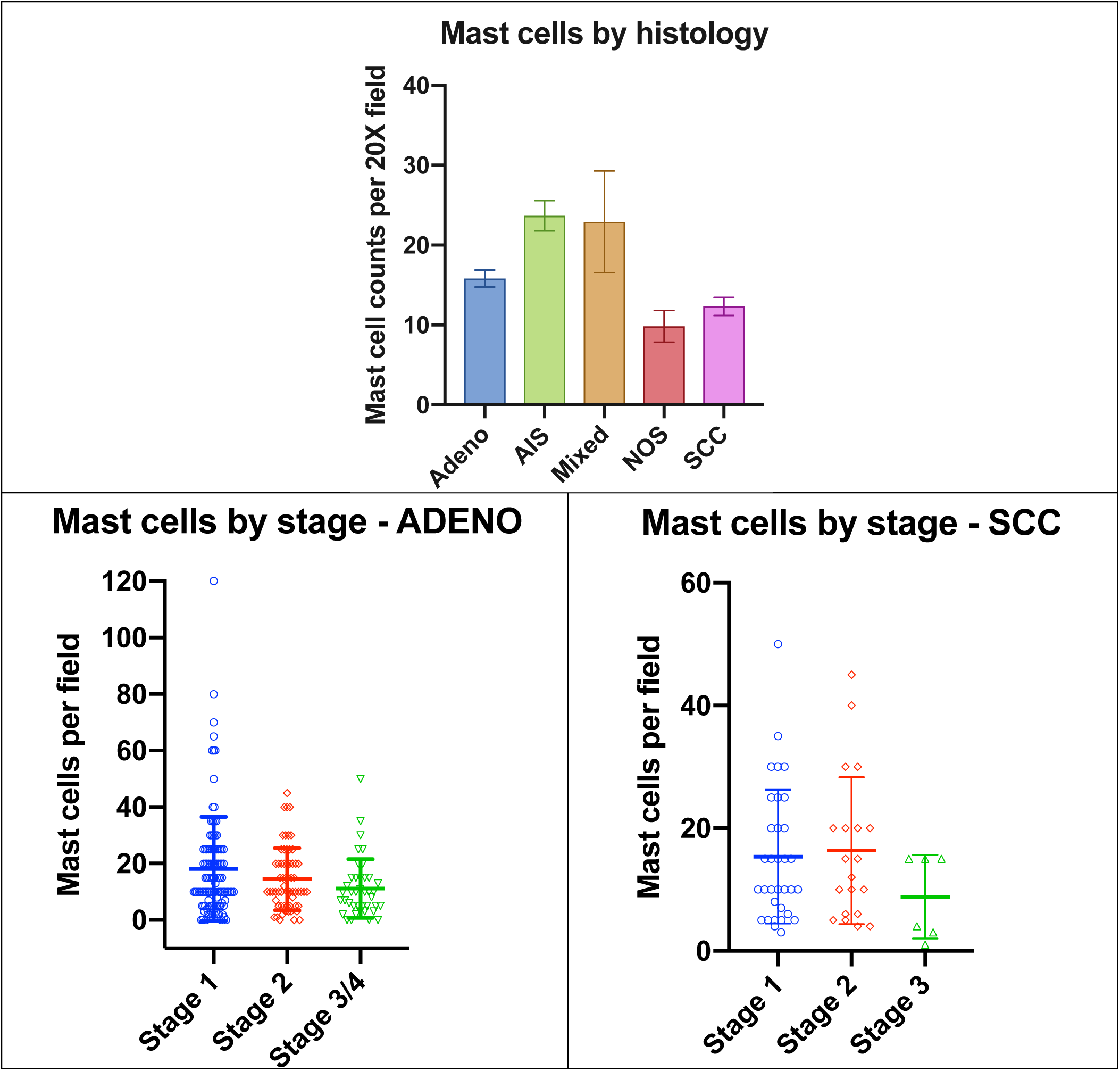
Mast cell counts across histologies, and across stages within adenocarcinoma and SCC. Mast cell counts across NSCLC histologies (top) on TMA; and across stage for adenocarcinoma (bottom left) and SCC (bottom right).

**Supplementary Figure 8:**
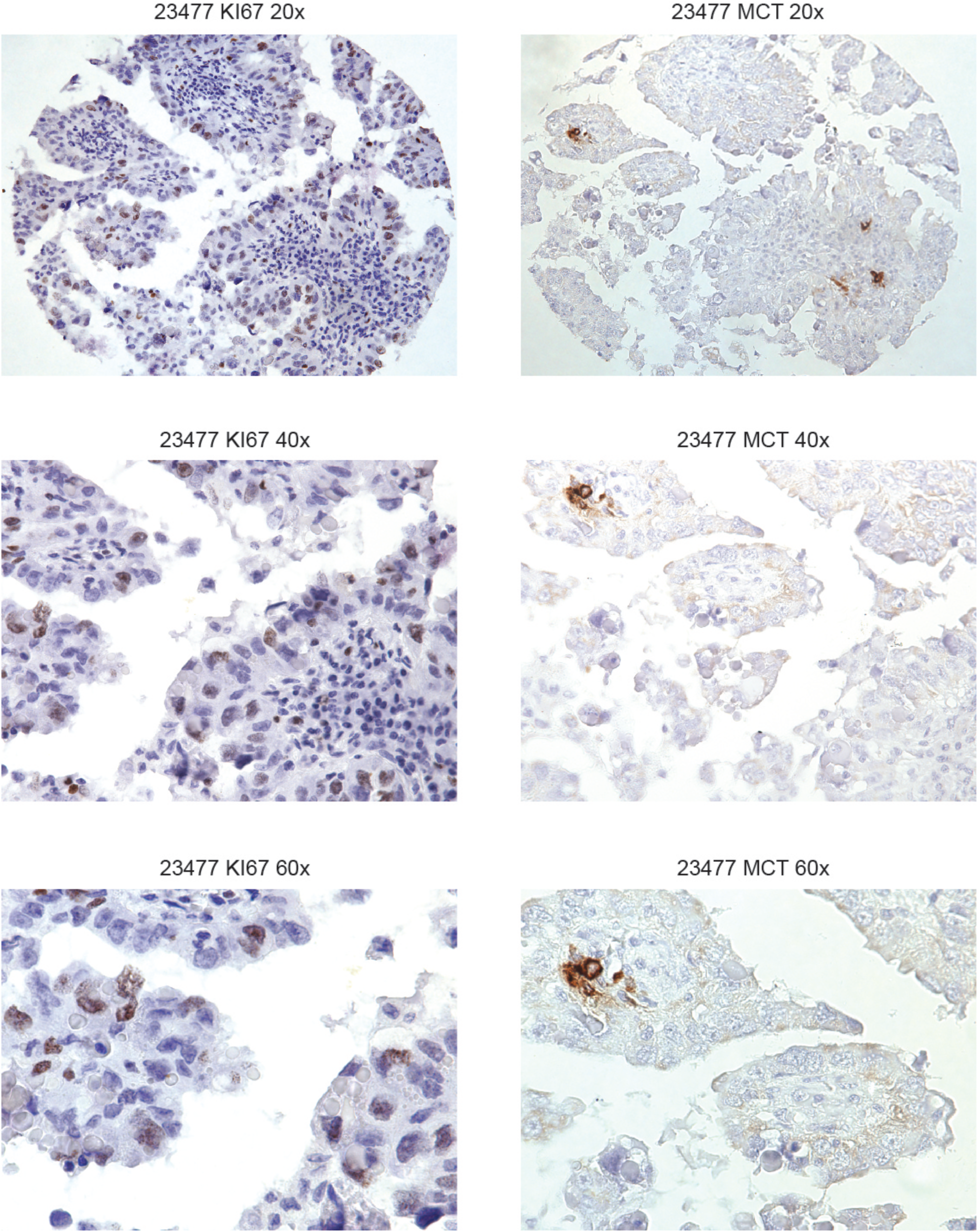
Mast cell tryptase and KI67 staining for a low-mast-cell infiltration adenocarcinoma.

**Supplementary Figure 9:**
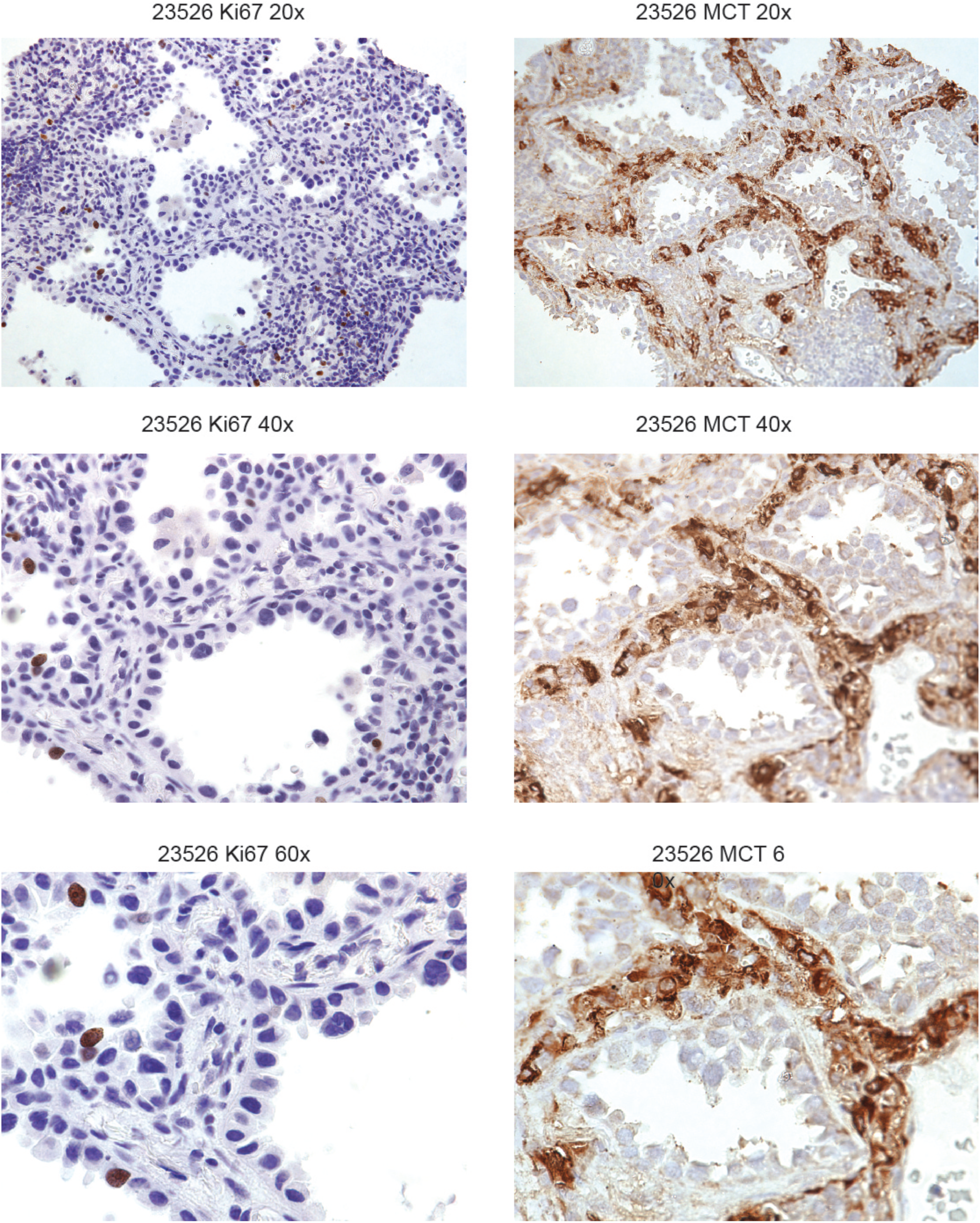
Mast cell tryptase and KI67 staining for a high-mast-cell infiltration adenocarcinoma.

**Supplementary Figure 10:**
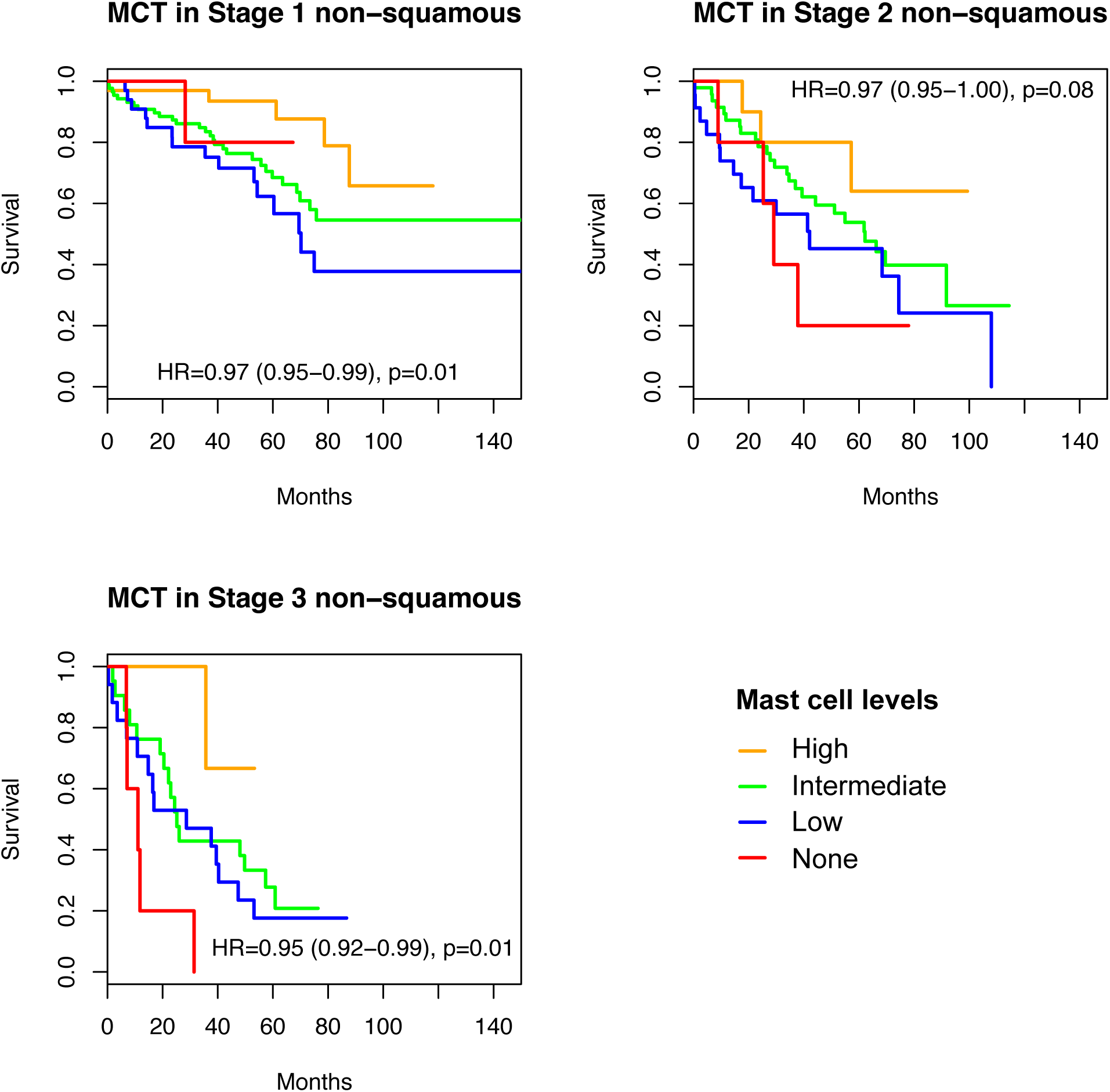
Kaplan-Meier analysis of Mast cell association with overall survival across stages of NSCLC.

